# Controlling Banana Bunchy Top Disease in Benin: crop protection strategies with socioeconomic perspectives

**DOI:** 10.1101/2025.07.24.666611

**Authors:** Renata Retkute, Martine Zandjanakou-Tachin, Bonaventure Aman Omondi, Ulrich Roland Agoi, Yaovi Marc Vodounou, Hermine Akofodji, Evrard Akpla, Laurence Dossou, Amélie Médénou, Alban Etchiha, Emilline Houménou, Alphonse Attadéou, Lava Kumar, John E. Thomas, Christopher A. Gilligan

## Abstract

**Summary:**

- This study combines satellite mapping, field surveillance, farmer surveys, and disease modelling to assess BBTV dynamics across Benin. It provides the first national-scale map of banana cultivation in Benin and identifies high-risk areas for targeted intervention.
- The study highlights the dominance of informal networks for exchange of planting material (∼80% of farmers), limited farmer awareness of BBTV (∼60% unaware), and low recognition of symptoms or transmission routes, all of which contribute to the spread and persistence of disease.
- Although women represented only a quarter of respondents, they demonstrated comparable levels of market orientation and access to planting materials and land. The findings call for spatially targeted, socially informed BBTV management strategies that integrate clean planting material systems, farmer education, and accessible control methods.
- BBTV was detected in 140 out of 747 fields surveyed, with the highest prevalence in the humid southern regions. No cases were found in the northern Sudanian zone, reflecting known agroecological patterns of disease distribution.
- Modelling of three control strategies - decapitation, uprooting, and injection - revealed injection as most effective nationally (70% reduction in disease incidence), while uprooting was most effective in vulnerable communities with high disease pressure.
- The study reveals an interconnected system in which socioeconomic vulnerability, informal seed networks, limited disease awareness, and spatial clustering of banana cultivation collectively drive BBTV dynamics in Benin, highlighting the need for epidemiologically effective, socially informed, and spatially targeted interventions.

**Societal Impact Statement:** Banana and plantain are critical for food security and income generation in West Africa, particularly for smallholder farmers in countries such as Benin. However, banana bunchy top virus (BBTV) poses a major threat to sustainable production. This study integrates high-resolution remote sensing, field epidemiology, mathematical modelling, and socioeconomic analysis to improve understanding of the spread of BBTV and to evaluate the effectiveness of different disease control strategies. Our findings reveal that BBTV risk is highest in southern Benin, where socioeconomic vulnerability is also greatest. Low disease awareness, limited adoption of effective control methods and informal exchange of planting material exacerbate this risk. By identifying priority areas and strategies tailored to local social, economic, and agroecological contexts, this research offers a roadmap for designing targeted, sustainable BBTV management programs. These insights can support smallholder resilience, reduce disease burden, and safeguard banana-based livelihoods across sub-Saharan Africa.

## 1 Introduction

Banana (*Musa* spp., encompassing banana and plantain) is a staple food crop across sub-Saharan Africa, playing a critical role in ensuring household food security and providing essential income for millions of smallholder farmers. In many regions, the crop serves as both a subsistence staple and a commercial commodity, contributing significantly to rural livelihoods and local economies. However, the sustainability of banana production is increasingly threatened by a range of biotic stresses, including viral diseases. Among these, banana bunchy top disease (BBTD), which is caused by banana bunchy top virus (BBTV: *Babuvirus musae*), has emerged as one of the most severe and persistent threats. The banana bunchy top virus is transmitted by the banana aphid (*Pentalonia nigronervosa*) and by the exchange of infected planting material. It can cause significant yield losses, even resulting in complete crop failure if unmanaged (Thomas, 2024).

In Benin, the first confirmed case of BBTV was reported in 2012 (Lokossou et al. 2012), raising concerns about its potential spread and long-term implications for national banana production. Despite the growing recognition of the threat, the epidemiology and spatial extent of BBTV in the country remain poorly understood, in part due to limited surveillance infrastructure and a lack of comprehensive data on host distribution and knowledge of planting material exchange practices.

To address these challenges, surveys were conducted in Benin from 2018 to 2021, aiming to characterize banana producers, production, knowledge and behaviour; to assess farmers’ varietal preferences and the diversity of banana cultivars in use; to map the geographic structure and extent of informal exchange networks for planting material; and to evaluate the current spatial distribution of BBTD across the country. By integrating survey data, spatial modelling, and high-resolution satellite imagery, the current study provides a foundational step toward understanding the landscape-level dynamics of BBTD in Benin and informing development of targeted management strategies.

## 2 Materials and Methods

### 2.1 Study area

A countrywide study was conducted in Benin (6-12°N; 0.40-3°E) during 2018-2021 (Figure 1A). Benin is located in the Dahomey gap, a savannah mosaic corridor that fragments the regional West African rainforest and is thought to have been induced by climate change during the Holocene (Salzmann and Hoelzmann 2005). Climatically, the country is subdivided in three main regions comprising semi-arid, sub-humid dry and sub-humid humid regions that are closely related to three ecological zone (Figure 1B). In sub-humid humid region, the climate is marked by the alternation of two rainy seasons (March-July; September) and two dry seasons (August; November-February) (Fink et al. 2017). The current study was carried out in the principal banana-producing regions of Benin, which are categorized into five major zones: the southeast, south, southwest, central, and northern areas of the country (Figure 1C). These zones were selected based on their high density of banana producers and the presence of pilot sites for Banana Bunchy Top Virus (BBTV) mitigation efforts.

**FIGURE 1.**
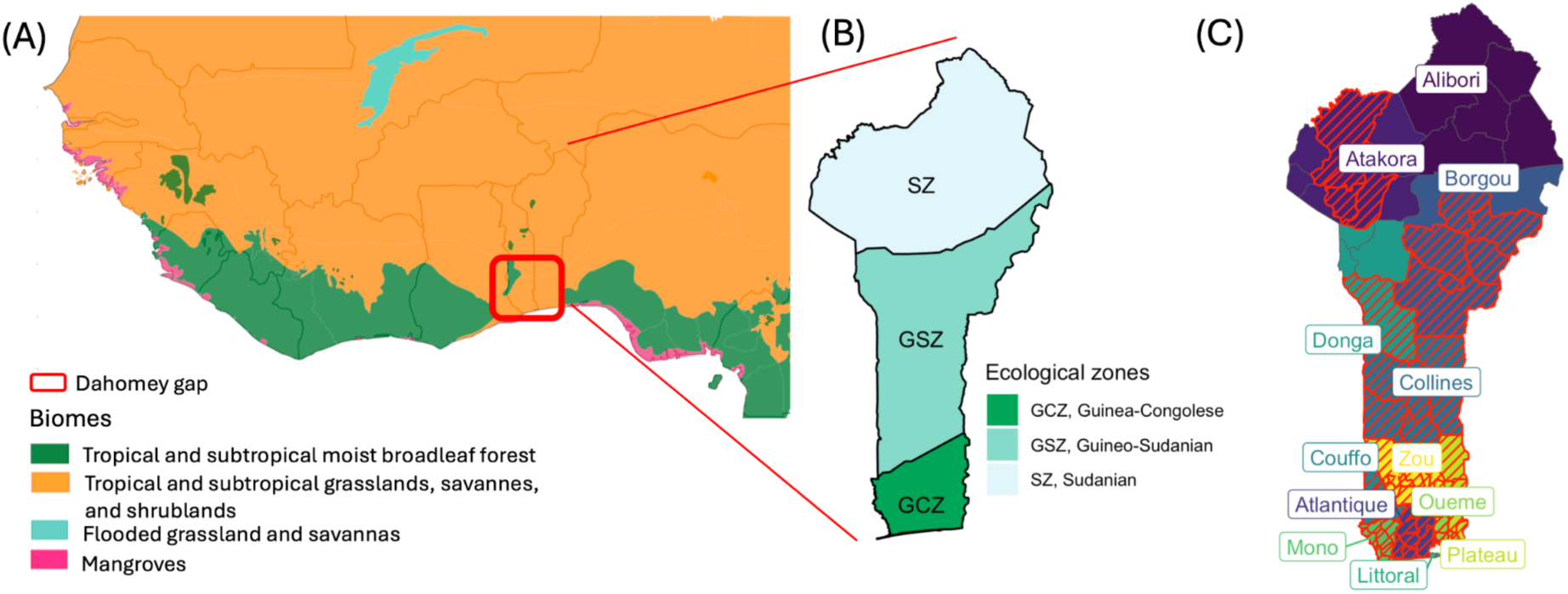
(A) Map showing biomes in West Africa (Olson et al., 2001). The Dahomey gap encircled. (B) Ecological zones in Benin: Sudanian Zone (SZ), Guineo-Sudanian Zone (GSZ), and Guineo-Congolese Zone (GCZ) (adopted from (Neuenschwander and Adomou 2017)). (C) Departments and communes in Benin, with surveyed communes highlighted as red hatched polygons.

### 2.2 Data collection

Farmers were selected using a non-probabilistic method (Trochim 2007). In each department in the selected zones, the main communes of banana production were identified from agricultural office databases. From each of the identified communes, two or three districts were selected and in each district two or three villages were surveyed. In the randomly selected villages, ten fields, separated by at least 1 km from each other, were surveyed. The field disease status was assessed based on characteristic symptoms such as morse code patterns of dark green streaks on the leaf petiole and lamina, and marginal leaf chlorosis (Hooks et al. 2008), using ten randomly selected mats. Farm coordinates were recorded using a GPS (Garmin, Etrex, Sommit HC). A questionnaire was used to record the farm history, cultivars grown and information about the farmer. Data on the efficacy of different disease management methods were collected from trained farmers participating in the project. Photographs of diseased plants presented to farmers are available in the Supplementary Materials (SI Figure A1).

### 2.3 Banana production map construction

We constructed a national-scale map of banana production in Benin by adapting the satellite-based classification framework of Retkute and Gilligan (2025b), integrating canopy height and NDVI data to distinguish banana cultivation from other vegetation. Using Google Earth Engine, we pre-processed Sentinel-2 images (August 2023–August 2024), excluding cloud-contaminated pixels and aggregated reflectance data into a median composite. Canopy height data from the Global Canopy Height Map (Tolan et al. 2024) were combined with annual mean NDVI to classify banana presence at 1 m resolution. This high-resolution binary classification was then aggregated to 250 m resolution to estimate banana coverage, after excluding unsuitable land comprising protected areas, water bodies, oil palm plantations, cashew crops, and built environments.

### 2.4 BBTV spread model

The BBTV spread model employs a spatially explicit susceptible-infected framework that also accounts for variations in banana densities and multiple pathogen entry pathways while incorporating localized increase in pathogen density at specific sites and facilitating virus transmission between grid cells (Retkute and Gilligan 2025b).

To parametrise the model using the survey data, we used the ABC-RF-rejection algorithm (Minter and Retkute 2019; Retkute and Gilligan 2025a). We employed two key summary statistics for the parameter estimation algorithm: (i) the sum of squared deviations between observed and simulated maximum radial expansion of BBTV across multiple time points; and (ii) the absolute difference between observed and simulated outbreak intensities.

### 2.5 BBTD management simulation

As a baseline for comparison, we simulated a hypothetical scenario in which no disease management interventions were implemented. This no-control scenario reflects conditions typical of resource-limited contexts, where farmer awareness, disease surveillance, and access to control measures remain low. We also evaluated three BBTV management strategies: decapitation, uprooting, and injection-based treatments. Each management strategy was modelled as a reduction in parameters governing BBTV spread. For uprooting, we assumed that the secondary transmission rate was reduced by 50%. For injection-based treatment, we assumed that the rate of infection build-up within a grid cell was reduced by 50%. To model the decapitation option, we reduced both the rates of primary and secondary transmission by 20%.

To identify the most vulnerable regions, we integrated simulation outputs with high-resolution spatial estimates of poverty across Benin, as provided by the Relative Wealth Index (RWI) from Chi et al. (2022). The RWI captures the relative economic standing of each location within the national context. We stratified locations based on two criteria: (i) the likelihood of BBTV spread, and (ii) socioeconomic vulnerability as measured by the RWI.

## 3 Results

### 3.1 Characteristics of banana producers, production, knowledge and behaviour

#### 3.1.1 Socio-demographic characteristics

A total of 627 individuals participated in the survey. Men comprised the majority of respondents, with women accounting for 23.1% of the sample (Figure 1A). Ethnic affiliation was diverse, with Fon being the largest ethnic group (42.4%), followed by Tori (26.8%) and Adja (24.4%), while Yoruba respondents comprised a minority (6.4%). Although the surveys were conducted in representative locations, certain ethnic groups - such as Mina, Bariba, Dendi, and Idasha - did not emerge among the dominant languages reported. The representation of women across the three major ethnic groups was relatively consistent, ranging from 21% to 26%, suggesting that the gender disparity observed is broadly distributed across ethnic groups. In terms of age distribution, respondents were primarily middle-aged: 45.6% were between 40 and 60 years old, and 38.7% were aged between 20 and 40 years. The proportion of female respondents in these age groups was also similar (26.7% in the 20-40 age group and 21.3% in the 40-60 age group), indicating that gender participation rates do not vary substantially across age cohorts. Most participants were married (over 90%), and educational attainment was generally low, with more than 60% having received no formal education.

Economic motivations were the dominant driver of banana cultivation: 81.4% of respondents reported growing bananas primarily for sale. Subsistence use was reported by only 9.4%, with an additional 9.2% cultivating bananas for both sale and household consumption. These figures were remarkably consistent across gender lines (82.0% of women and 81.3% of men engaged in banana production for sale) and age groups (82.3% among those aged 20-40 and 81.8% among those aged 40-60).

#### 3.1.2 Production characteristics

Survey data were collected from six of Benin’s twelve administrative departments, with a notable concentration of responses from the southern regions (Figure 1B). The largest share of respondents originated from Ouémé (36.0%), followed by Mono (21.4%), Plateau (16.3%), Atlantique (14.3%), Zou (7.0%), and Couffo (5.0%). Across all surveyed departments, agriculture was overwhelmingly reported as the primary livelihood activity, with 95% of respondents identifying it as the main economic pursuit of their household. Small variations in subsistence-oriented banana production were observed, ranging from 0% in Couffo to modest levels in Zou and Plateau (<4%) and reaching approximately 10% in Mono, Atlantique, and Ouémé. Banana cultivation was primarily conducted on small-scale plots. Only 7.3% of respondents reported owning plantations larger than 1,000 m² (equivalent to >0.1 ha). Approximately one-third of plantations measured less than 100 m², and 45.6% fell between 100–500 m². Notably, larger plantations (>1,000 m²) were concentrated in Couffo (26.1%), Mono (28.2%), and Ouémé (23.9%). Among owners of larger plantations, 21.7% were women - closely mirroring the overall gender representation in the survey - indicating no disproportionate access by gender to larger landholdings in banana farming compared with the overall population of banana growers. In terms of land tenure in banana cultivation, nearly half of the respondents (48.1%) had owned their plantations for 1-5 years, and one-third had done so for 6-10 years. Long-term landholding was uncommon; only ∼5% had cultivated bananas on the same plot for more than 20 years. Of these long-established farms, 50% had areas between 100–500 m², and 90% were owned by men. Only 1.4% of respondents had established their plantation within the year preceding the survey.

The majority of respondents (79.6%) reported sourcing banana planting material from neighbours, with smaller proportions using material from their own fields (14.2%) or other sources (6.3%). The widespread reliance on neighbour-sourced planting material was broadly consistent across commercial (80.8%) and subsistence (71.2%) producers, and between female (82,7%) and male (78,6%) respondents.

#### 3.1.3 BBTD awareness, detection, and management practices

Banana bunchy top disease was detected at 47.4% of households growing bananas primarily for subsistence, compared with 27.5% of those cultivating bananas for commercial purposes (Figure 2C). Despite the observed presence of the disease, overall awareness was limited: approximately 60% of respondents reported having no knowledge of BBTV. Among farmers in whose plantations BBTD was detected, 70% had at least heard of the virus, yet only 40% reported a clear understanding or awareness of the disease. In stark contrast, just 20% of farmers in BBTD-negative households had any knowledge or awareness of the virus. Among those familiar with BBTV, key sources of information included the National University of Agriculture through the “project BBTV” initiative (19%) and informal community networks (21.4%, primarily neighbours). Recognition of BBTD-associated symptoms remains low. Only 11.3% of respondents claimed they could identify BBTD symptoms in the field, and 16.1% could do so from photographs. As a result, it is unsurprising that 62.3% of farmers with BBTV-infected plantations reported they would not recognize the disease in situ. Proactive monitoring efforts were limited: only 20% of farmers reported actively inspecting their fields for BBTD symptoms, and just 11.3% were engaged in targeted scouting for infected plants. Notably, 81.6% of those practicing active surveillance managed plantations of 500 m² or less, suggesting that smaller-scale farmers may be more attentive or capable of field-level inspection, perhaps due to greater manageability. Understanding of BBTV infection and transmission was similarly low. While 59.1% of respondents did not know the cause of infection, 90% were unfamiliar with transmission pathways. Lack of maintenance (62.8%), contaminated planting material (24.2%), and pests such as insects or fungi (12.9%) were cited as possible causes. Suggested transmission routes included aphids (55.3%), and infected discharge (42.8%). Some of the farmers were aware of the presence of other banana diseases in their plantations: 7.6% mentioned Fusarium wilt, 7.0% cited Sigatoka, and 3.8% identified Banana Xanthomonas wilt.

**FIGURE 2.**
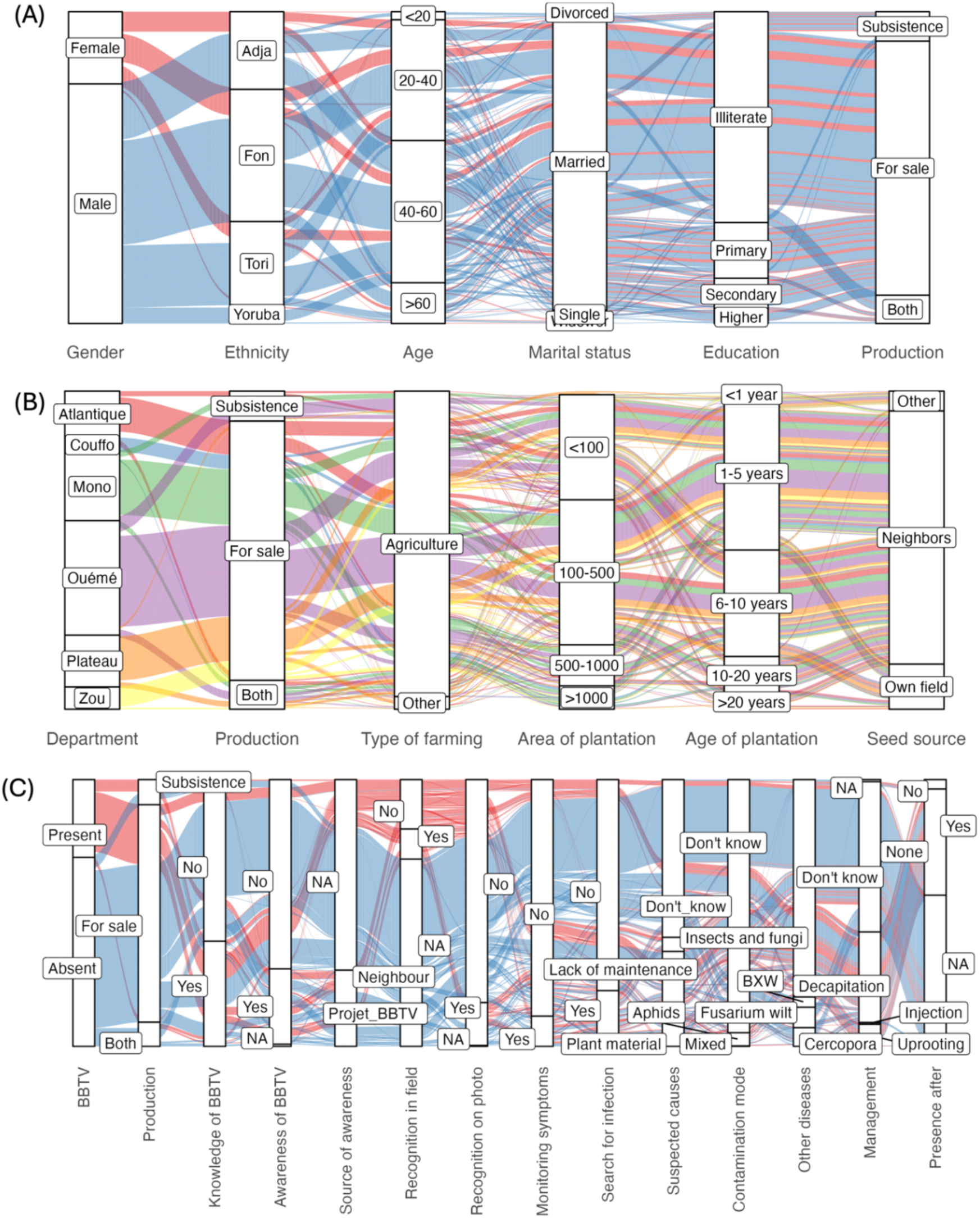
Sankey plots derived from farmer surveys showing: (A) relationships between socio-economic factors describing the producers; (B) relationships between agronomic factors describing the production; and (C) relationship between status of BBTV on farms and knowledge/behaviour of producers concerning the pathogen.

When queried about disease management practices specific to BBTV, more than half (56.3%) reported taking no action. Among those who engaged in any management, the most common responses were decapitation of symptomatic plants (33.9%), uprooting (8.2%), and, rarely, injection-based treatments (0.6%). According to farmer perceptions, 39.8% of respondents believed that BBTV may continue to persist following the intervention.

### 3.2 Cultivar selection for banana production

The selection of banana cultivars by farmers is shaped by a complex interplay of factors, including local availability, agronomic performance (e.g., yield potential, resistance to biotic and abiotic stresses), and market or consumer preferences such as taste and commercial value. Across all survey responses, a total of 21 distinct-cultivars were documented: *Agba, Ahlor, Alogli, Alokpoe, Avlan, Aminninè, Aoukokoé, Chinoise, Dankokoé, Danvlan, Gangnikokoé, Gbakokoé, Gbogui, Gounkokoé, Honkè, Oliri, Planta, Sokokoué, Sotoumon, Tchèkètè,* and *Tchon* and undefined plantain cultivars. Some characteristics of the cultivars are shown in the Supplementary Materials Table A1.

A majority of respondents (54.5%) reported growing a single cultivar, a practice dominant across all surveyed departments (Figure 3A), and particularly prevalent in Plateau (68.6%) and Mono (58.2%). Cultivation of two cultivars was reported by 35.95% of respondents, while fewer than 10% grew more than two. Plantain types were most frequently cultivated, mentioned by 49.1% of respondents, with widespread presence across all departments (Figure 3B). *Planta* and *Sotoumon* were also commonly reported, whereas the remaining 19 cultivars appeared in only one or two departments, suggesting either limited geographic distribution or potential gaps in varietal identification among farmers.

**FIGURE 3.**
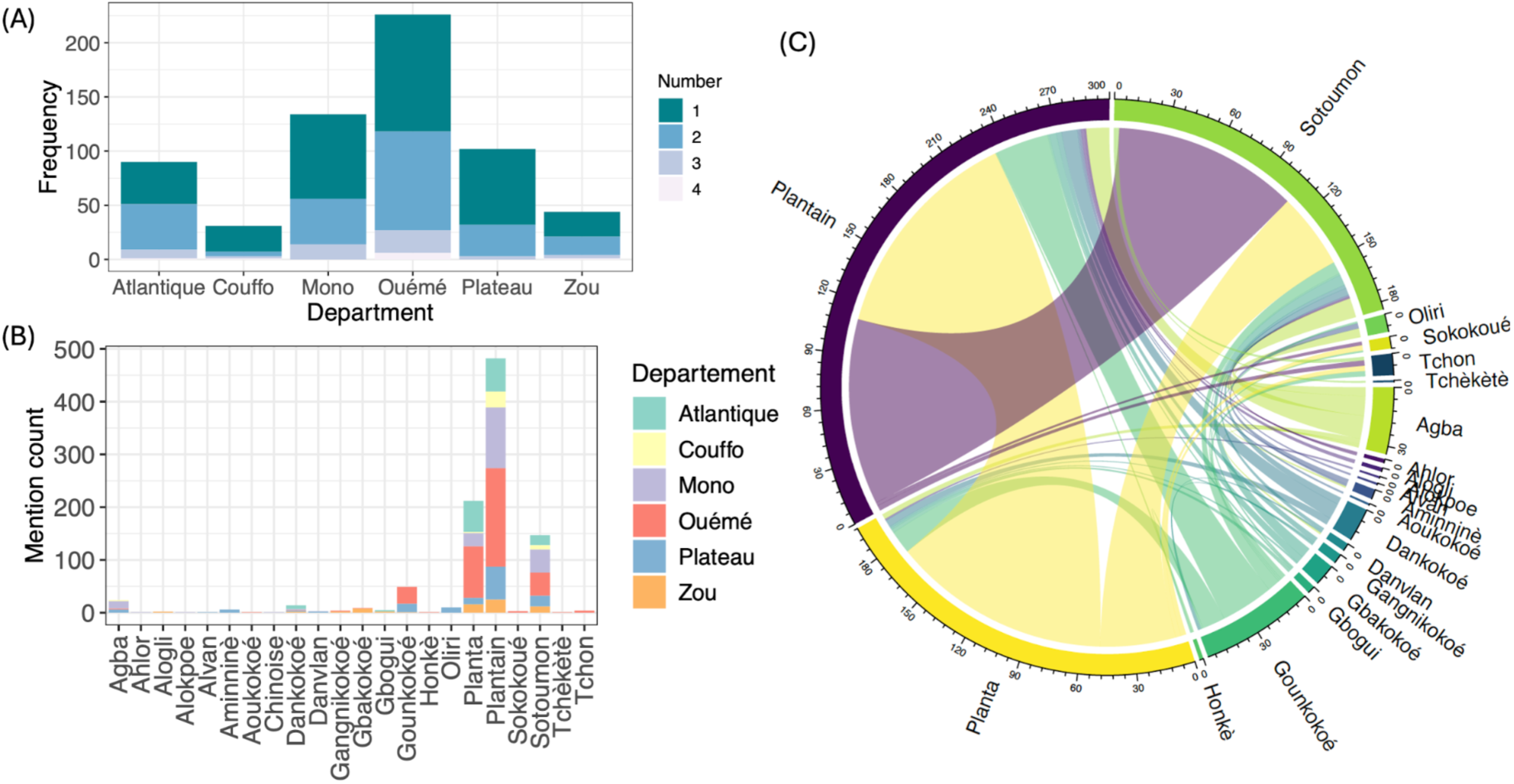
Analysis of banana and plantain cultivar information from farmer survey. (A) Frequency of cultivation of a single cultivar (1) and multiple cultivar (two or more) mentioned in the same response. (B) Frequency of individual variety mentions across all survey responses stratified by department. (C) Patterns of cultivar co-occurrence reported within the same survey response, highlighting frequency of cultivar combinations. Numbers outside the circle indicate the count of mentions in responses listing multiple cultivars.

To understand patterns of varietal co-cultivation better, we analyzed co-occurrence of cultivars within individual survey responses (Figure 3C). The most frequent combinations were “plantain” with *Planta* (29.0%) and “plantain” with *Sotoumon* (24.5%). Notably, 12% of respondents reported growing *Gounkokoé* alongside one or more of the three most common cultivars.

### 3.3 Geographic patterns of banana and plantain seed exchange networks

Information on the geographic origin of planting material was provided by 66.1% of respondents. These data were analyzed at the level of second-order administrative divisions (communes) in Benin, allowing for the reconstruction of sourcing networks for planting material across 18 communes (Figure 4). The complexity and geographic extent of these networks varied substantially among locations.

**FIGURE 4.**
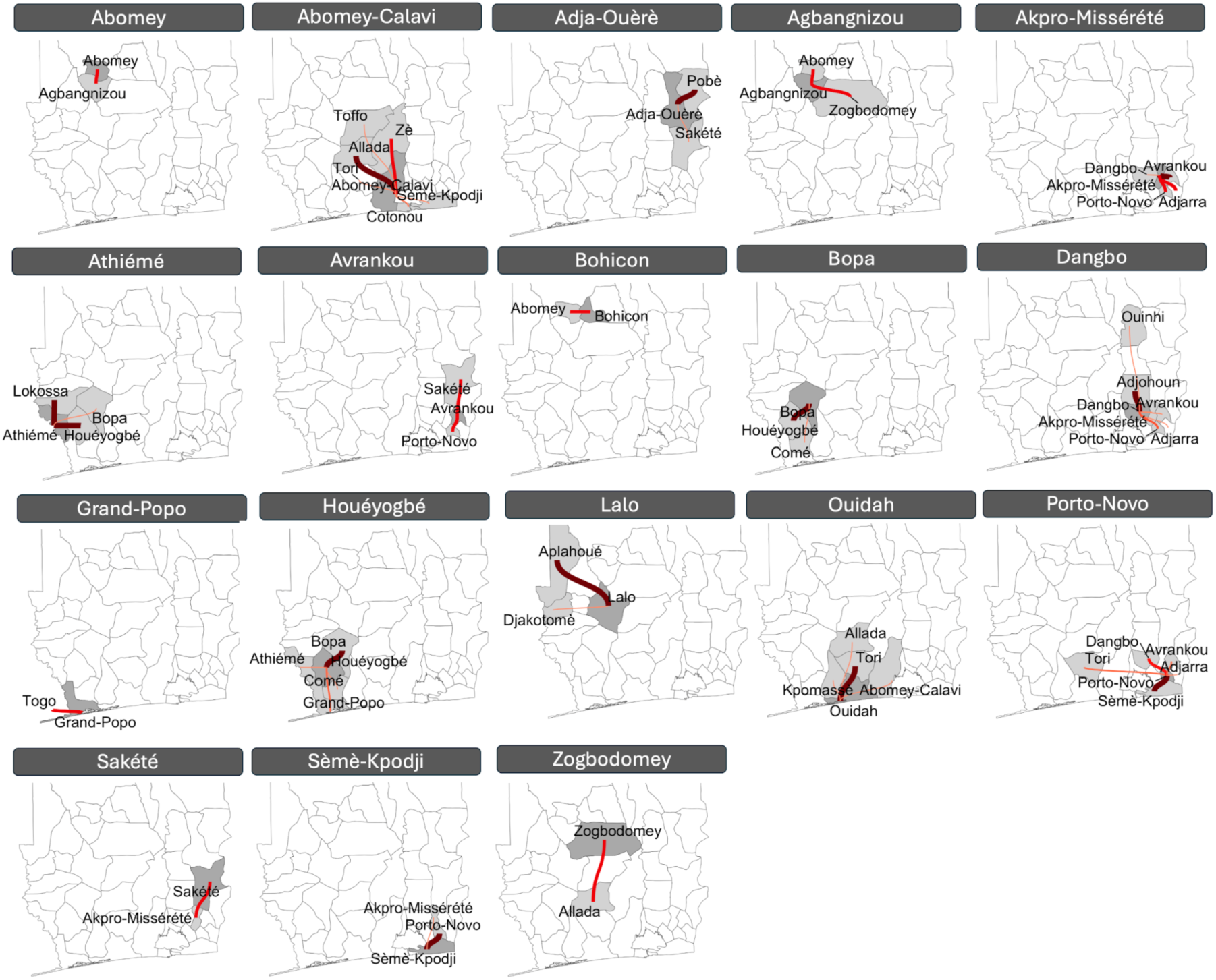
Network for exchange of planting material amongst communes in southern Benin. Dark grey shows receiving communes and light grey shows donor communes for sourcing planting material. Intensity and width of lines indicate relative frequency of reported movement for each target commune.

The most intricate sourcing networks were observed in Abomey-Calavi, Porto-Novo, and Dangbo. In Abomey-Calavi, farmers reported sourcing planting material from seven different communes, with Allada emerging as the most frequent source. Additional sourcing occurred from neighbouring communes such as Tori and Zè, as well as coastal communes including Sèmè-Kpodji and Cotonou. Similarly, Dangbo respondents reported sourcing planting material from seven communes, primarily from adjacent Adjohoun, with more distant sourcing from Ouinhi, approximately 60 km away. In Porto-Novo, six source communes were identified, with 56% of the planting material coming from nearby Sèmè-Kpodji and Avrankou. The most distant source was Tori, located roughly 80 km away. Tori also served as a notable source for farmers in Ouidah, despite not being a neighbouring commune. In contrast, respondents in Houéyogbé sourced planting material from four of their five adjacent communes. This preference for neighbouring sources was also observed in several other communes, such as Abomey and Bohicon (one neighbouring commune) and Adja-Ouèrè, Agbangnizoun, Athiémé, Avrankou, Bopa, Ouidah, and Sakété (two neighbouring communes).

Exceptions to the localized pattern included Lalo and Zogbodomey, where farmers reported sourcing planting material exclusively from non-neighbouring, more distant communes. Notably, two respondents in Grand-Popo cited Togo as their source, indicating exchange across a national boundary.

### 3.4 BBTV surveillance results

Between October 2018 and June 2021, BBTD was detected in 140 of the 747 fields surveyed across Benin (Figure 5). Notably, no BBTV cases were identified in any of the 37 surveys conducted within the Sudanian ecological zone (SI Figure A2A), which included surveys from the Atakora Department in 2018 and the Borgou Department in 2021 (SI Figure A2B). The prevalence of BBTV was comparable across the other two major ecological zones: 25.3% of surveys reported BBTV presence in the Guineo-Sudanian zone, and 18.3% in the Guineo-Congolese zone. Surveillance efforts were most intensive in 2019, accounting for 51.9% of all surveys, with similar levels of activity (110–140 surveys) in the remaining years (Figure 5).

**FIGURE 5.**
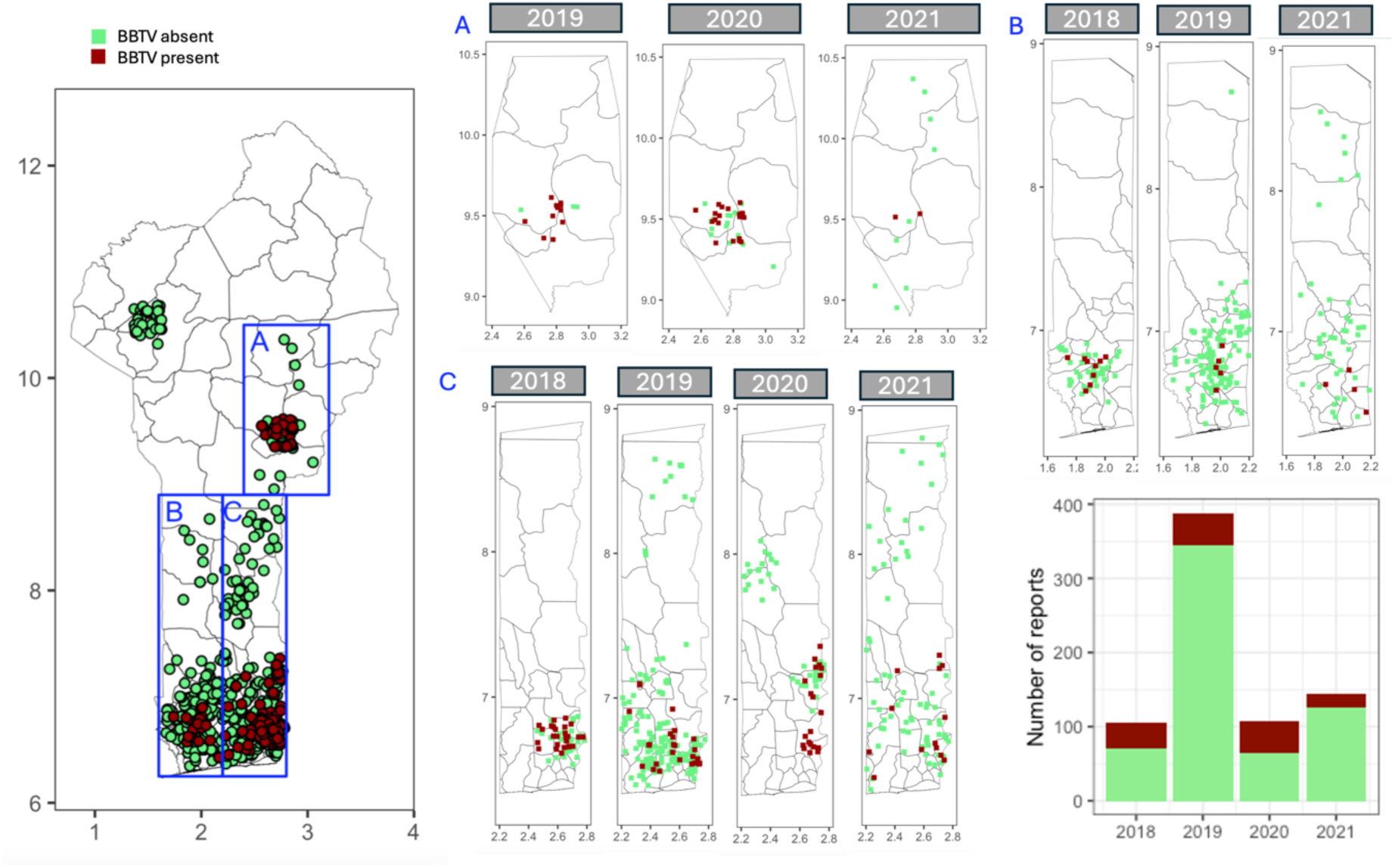
BBTV surveillance: temporally aggregated (left) and aggregated at a grid cell level (250 m x 250 m) for three blocks (A, B and C, blue rectangles). The lower right figure shows the total number of BBTV absence (light green) and presence (dark red) reports per year.

To characterize spatial patterns of disease emergence, we aggregated survey data to a 250-meter grid resolution and identified three distinct clusters of active BBTV transmission (Figure 6). Only Cluster C was monitored consistently over four consecutive years (2018–2021). The maximum radial spread observed was 32 km after three years in Cluster A, 64 km after four years in Cluster B, and 85 km in Cluster C. The proportion of infected grid cells within each cluster was 0.561 for Cluster A, 0.088 for Cluster B, and 0.229 for Cluster C (SI Table A2).

**FIGURE 6.**
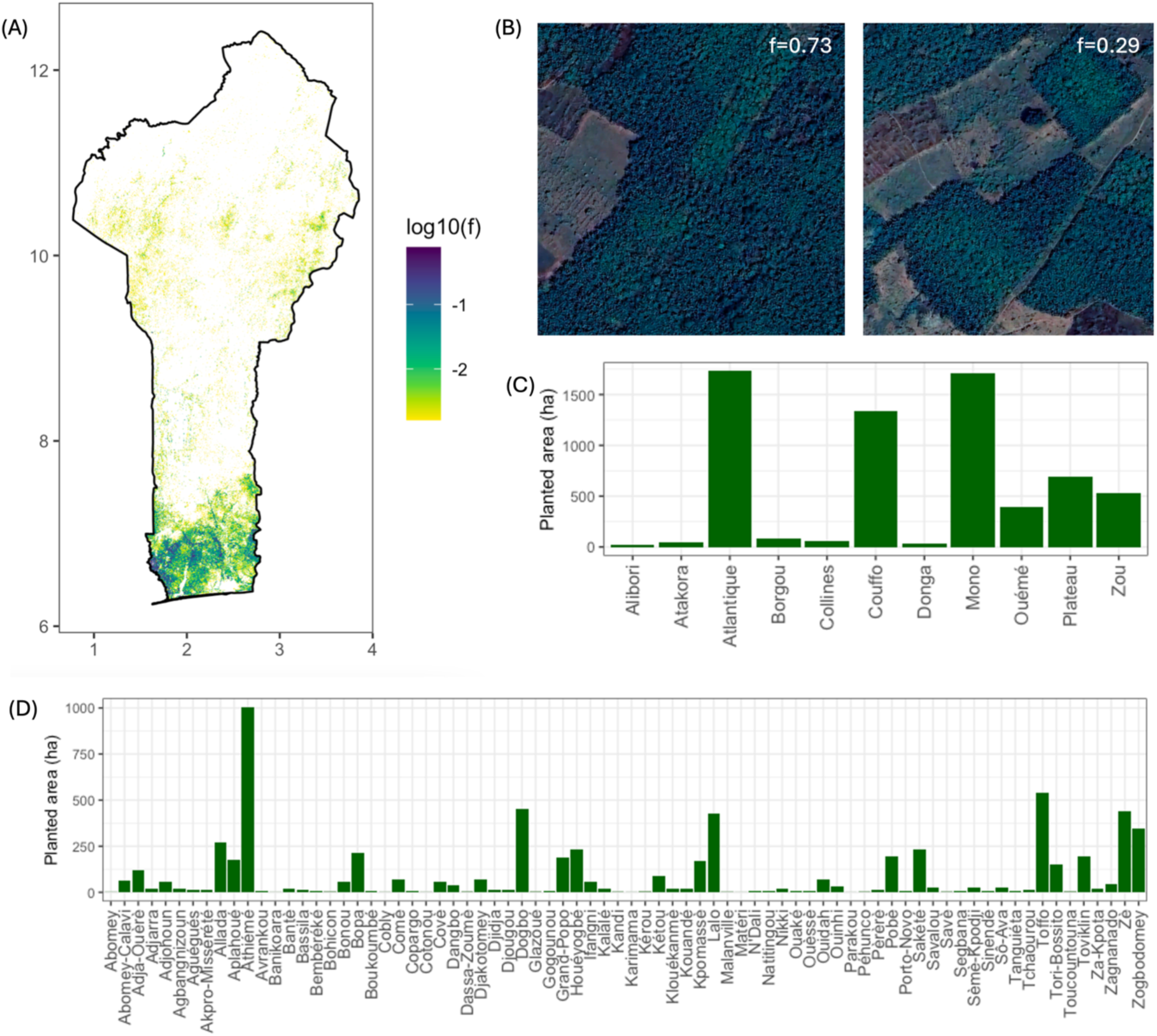
(A) Distribution map for banana cultivation in Benin, estimated using a combination of NDVI and canopy height data derived from high-resolution satellite imagery at 1 m resolution. Shading shows log10 fraction of a 250 m x 250 m grid cell planted with bananas. (B) Examples of grid cells with high (*f =* 0.73) and medium (*f =* 0.29) density of bananas. Satellite images in (B) obtained from Google Earth Engine (Imagery © 2024 Airbus, CNES/Airbus, Landsat/Copernicus, Maxar Technologies). (C) Estimated area planted by bananas in Departments. (D) Estimated area planted by bananas in Communes.

### 3.5 National-scale map of banana production in Benin

Using high-resolution satellite remote sensing data and a refined classification framework incorporating canopy height and vegetation greenness (NDVI) thresholds, we generated a spatially explicit map of banana cultivation across Benin at a 250 m spatial resolution (Figure 6A). This integrative approach leverages structural (canopy height) and phenological (NDVI) characteristics to discriminate banana production areas from other vegetation types, including perennial crops, natural forests, and annual field crops. Our analysis estimates the total banana-harvested area in Benin to be approximately 6,663 hectares. We found no correlation between our estimates aggregated at 10 km resolution and banana cultivation estimates from the SPAM 2020 dataset (Yu et al. 2020) (SI Figure A3).

The resulting banana presence map reveals a highly heterogeneous spatial distribution of banana and plantain cultivation across Benin, with a pronounced concentration in the southern regions of the country. Visual inspection of 20 grid cells sampled at various ranges of cover demonstrated strong concordance between satellite-based estimates and observable land use (SI Figure A4). Two such examples are shown in Figure 5B. Estimates of cropped area were further computed at the department (Figure 6C) and commune levels (Figure 6D), based on spatial aggregation within each administrative boundary. The highest banana densities were recorded in the southern departments of Atlantique (1,731 ha), Mono (1,704 ha), Couffo (1,343 ha), Plateau (697 ha), Zou (531 ha), and Ouémé (398 ha). Together, these six departments account for 96.1% of the total area under banana and plantain cultivation in Benin. At the commune level, the five communes with the largest cultivated areas were Athiémé (1,002 ha), Toffo (541 ha), Dogbo (455 ha), Zè (440 ha), and Lalo (428 ha), together comprising 43% of the total area under banana cultivation in Benin.

### 3.6 Parameterisation of BBTV spread model

Model parameterization was based on BBTV surveillance data collected between 2018 and 2021 (Section 3.4). Outbreaks in Blocks A, B, and C (Figure 5) were modelled as independent events - an assumption supported by the survey data presented in Section 3.3, particularly the wide spatial distribution of clean planting material. Parameter estimation was carried out separately for each block. In the first stage of the ABC-RF algorithm, 10,000 parameter sets were sampled and simulated per block, retaining only those that closely matched the observed spread patterns. This process yielded 14, 22, and 17 accepted sets for Blocks A, B, and C, respectively. A Random Forest model was then trained on these accepted particles, and 10⁷ new parameter sets were generated and evaluated, with further simulations performed only for those with acceptance probabilities above 0.3. After this second round of parameter estimation, 215, 257, and 226 accepted sets were obtained for Blocks A, B, and C, respectively. Summary statistics of the accepted particles from stage two, along with the outcome of a representative posterior simulation of BBTV spread, are shown in Figure 7. The resulting block-specific posterior distributions were subsequently combined to construct an overall posterior distribution (SI Figure A5). To account for the implementation of disease mitigation measures by a subset of farmers, the fitted parameter estimates were adjusted using management data derived from farmer surveys. Further details are provided in Supplementary Information, Section A7.

**FIGURE 7.**
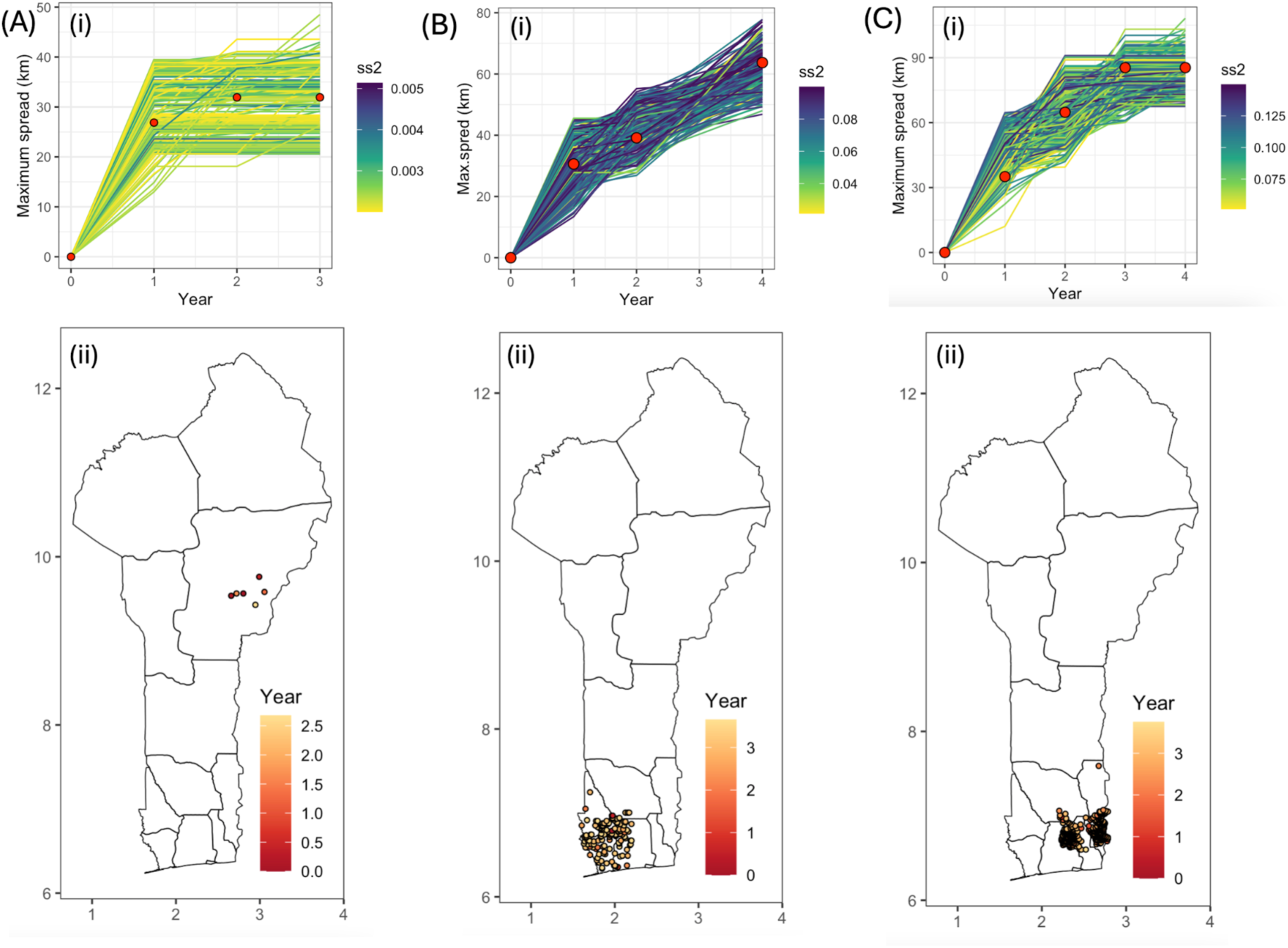
Parameterisation of the BBTV spread model for Blocks A, B, and C. Panels (A), (B), and (C) each display (i) summary statistics derived from posterior parameter sets obtained after two-stage ABC-RF estimation (top row), and (ii) simulated BBTV spread based on a representative posterior parameter set for the respective block (bottom row). Red points on top row indicate the observed spatial extent of BBTV spread.

### 3.7 Assessing the impact of BBTV management strategies

To evaluate the potential spread of Banana Bunchy Top Disease (BBTD) in the absence of control efforts, we simulated a hypothetical scenario in which no disease management interventions were implemented. Simulations initiated outbreaks in each of the three spatial blocks, with the index infection seeded in randomly selected grid cells that tested BBTD-positive in baseline surveys - 2018 for Blocks B and C, and 2019 for Block A. Simulations were run over a six-year period, covering the timeframe from initial detection to early 2025. Disease risk was expressed as the proportion of simulations in which a given grid cell became infected, denoted as BBTV risk (*R*). Results indicated that BBTV risk was highest in the southern regions of the study area, which corresponds with the primary banana-producing zones (Figure 8A). In contrast, the Relative Wealth Index (RWI) displayed high spatial variability, with wealth concentrated around urban centres (Figure 8B).

**FIGURE 8.**
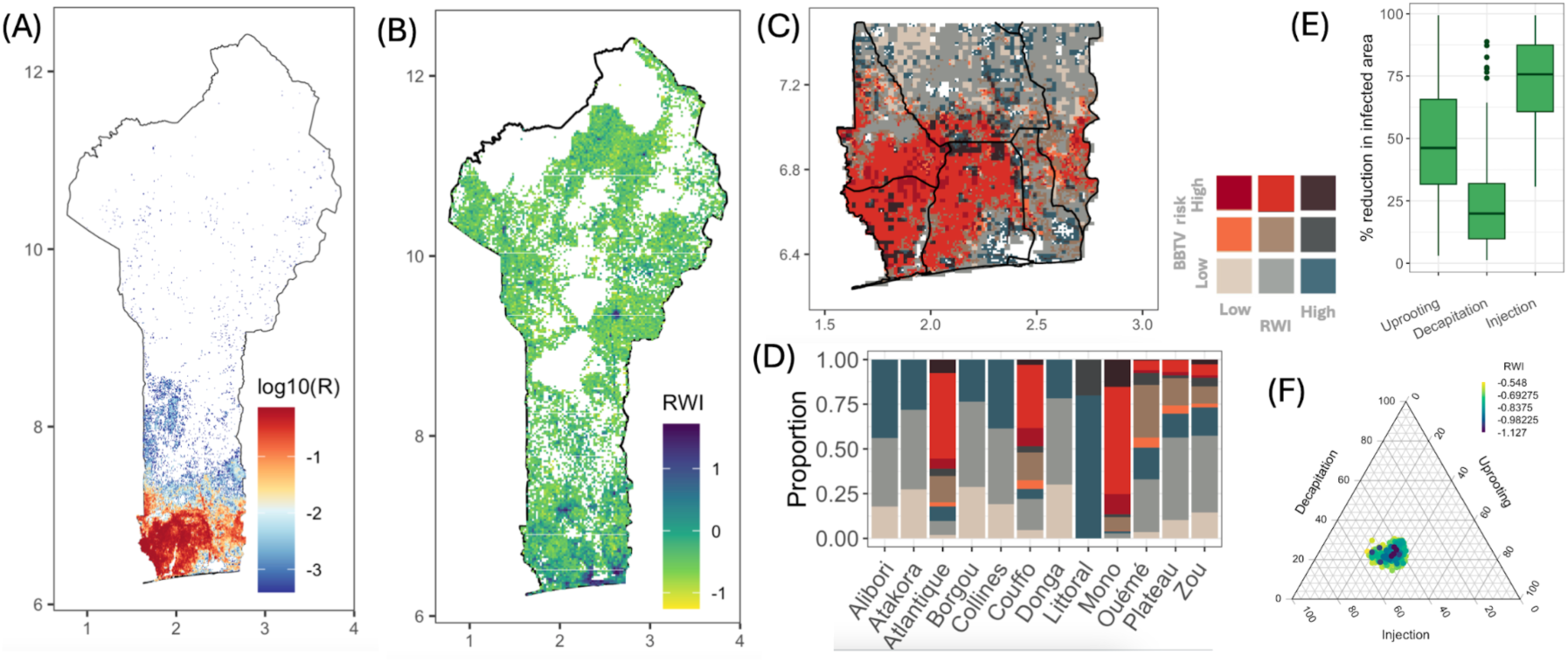
(A) Prospective spread of BBTD under a hypothetical scenario with no disease management interventions. The BBTV risk (*R*) is defined as the proportion of simulations in which BBTV was present in a grid cell. (B) Spatial distribution of Relative Wealth Index (RWI) (Chi et al. 2022). (C) Spatial distribution of simulated BBTV risk and RWI in the southern region. BBTV risk and RWI were each divided into three classes: Low, Medium and High. (D). Proportion of combined BBTV risk and RWI classes in each Department. (E) Reduction in infected area due to three control strategies (decapitation, uprooting and injection) at a national scale. (F) Effect of control strategies for grid cells with high BBTV risk and low RWI.

By integrating modelled BBTV risk with the RWI, we identified areas where high disease pressure coincides with greater vulnerability (Figure 8C). Grid cells are classified into low (L), medium (M), or high (H) categories based on BBTV risk and RWI values (SI Table A4). The departments of Mono, Atlantique, and Couffo were found to be most affected, with 71.2%, 54.0%, and 45.7% of their areas, respectively, falling into high BBTV risk and low-to-medium RWI categories (Figure 8D). Three additional departments (Ouémé, Plateau, and Zou) had smaller proportions (<10%) of similarly affected areas. In total, approximately 9% of Benin’s land area was identified as both highly exposed to BBTV and socioeconomically disadvantaged. Five communes (Athiémé, Bopa, Houéyogbé, Comè, and Grand-Popo) were situated at the intersection of high epidemiological risk, socio-economic vulnerability (Figure 8C), as well as being sources of planting material exchange networks in southern Benin (Figure 4). Within these communes, approximately 73% of the land area was classified as both high-risk for BBTV and within the low-to-medium RWI bracket, underscoring their significance as epidemiological and socio-economic stepping stones for disease spread.

To evaluate the potential effectiveness of BBTV control strategies, we ran simulations incorporating three different intervention scenarios: injection-based treatment, uprooting, and decapitation (SI Table A3). When comparing simulated control strategies, injection-based treatments demonstrated the highest efficacy at the national scale, achieving an average reduction in infected area of 73.5% (95% CI: 43%–95%) relative to the no-control baseline (Figure 8E). Uprooting provided moderate effectiveness, with a mean reduction of 48.3% (95% CI: 17%–87%), while decapitation was the least effective strategy, yielding a 24.7% average reduction (95% CI: 2.5%–65%). To assess which interventions may yield the greatest impact in communities with the greatest need, we focused on locations with both high BBTV risk and the lowest RWI values. Within these high-priority grid cells, uprooting emerged as the most effective local intervention, reducing BBTV risk by 70–85% after application for six years (Figure 8F-E).

## 4 Discussion

This study provides critical insights into the complex dynamics of BBTV in Benin. The virus poses a major threat to banana and plantain crops, which are essential for food security and income generation among smallholder farmers throughout sub-Saharan Africa. By integrating mathematical modelling with field surveillance and socioeconomic data, we have quantified the impact of different control strategies while also identifying priority areas where interventions could be most effective, considering both epidemiological risk and local socioeconomic vulnerability. Such a comprehensive, data-driven framework is valuable for designing targeted, sustainable management programs that support smallholder resilience and protect these critical food systems against ongoing and future disease challenges.

A key insight from the household survey was the strong economic incentive driving banana cultivation. The majority of respondents reported growing bananas primarily for market sale, underscoring the crop’s dual function in supporting both food security and income generation. The market price for a single bunch ranges from USD 2.50 to USD 20.00 between January and May, depending on the cultivar and seasonal availability (personal communication, M.Z.T.). This price range offers smallholder farmers valuable opportunities to generate income and strengthen their financial resilience. In coastal West African countries like Benin, agricultural landholdings are typically fragmented and small in scale; over half of farms in Benin are classified as very small (<0.15 ha) (Pérez-Hoyos et al. 2017). Previous estimates place banana field sizes in Benin between 0.01 ha and 1 ha, with an average of approximately 0.2 ha (Chabi et al. 2018). Our findings align with this range, revealing that banana cultivation remains predominantly small-scale, with most plots measuring less than 0.05 ha, yet it remains a deeply integrated component of rural livelihood systems.

The study highlighted notable cultivar diversity within Benin’s banana and plantain production systems, documenting 21 distinct cultivars in addition to plantains. Despite this diversity, a majority of farmers cultivated only a single cultivar, with plantains dominating across all surveyed regions. Among the most widely adopted cultivars were *Sotoumon* and *Planta,* while the remaining varieties were typically localized to one or two departments. These patterns mirror regional findings; for instance, Vodournou et al. (2022) reported that Plantain (*Alogo*) and *Planta* were the most frequently cultivated varieties in Benin. Farmers’ varietal choices are shaped by several interrelated factors, including local availability (Gold et al. 2002), taste (Ahouangninou et al. 2021), agronomic traits such as yield and resilience (Chabi et al. 2023), and market preferences (Madalla et al. 2023). For example, (Abiola et al. 2024) noted that challenges in marketing plantain, particularly low selling prices, influenced farmer decisions. These findings highlight the critical need to establish clean planting material systems that are not only technically robust but also aligned with farmers’ cultivar preferences and economic constraints. At the same time, while cultivar diversity provides agronomic and market benefits, it may pose challenges for effective BBTV detection and control, as symptom expression can vary significantly across banana varieties.

The sustainability and vulnerability of banana production are strongly shaped by how planting materials are sourced. In Benin, there are no certified suppliers of banana planting material, with the exception of the Institut de Recherches Agricoles du Bénin (IRAB), which maintains a dedicated banana research program. Additionally, the National University of Agriculture has supported pilot sites by establishing demonstration farms and distributing planting material.

The lack of access to clean planting materials has previously been identified as a major constraint to banana production in Benin (Ahouangninou et al. 2021). Approximately 80% of farmers reported acquiring planting material from neighbours, underscoring the dominance of informal, community-driven planting material exchange systems. These networks are often rooted in familial and social ties within and across villages, which serve as the foundation for their persistence and functionality (Nduwimana et al. 2022). Seed sourcing from neighbouring farmers is primarily driven by economic considerations, specifically, the availability of high-demand or high-value varieties and the reduced transportation costs associated with local proximity. While these decentralized systems are critical for ensuring access and affordability, especially for resource-limited farmers, they also represent a significant pathway for the transmission of BBTV, primarily through the exchange of infected suckers. For instance, gardens planted with suckers obtained from neighbours have been found to be 53 times more likely to harbour BBTV compared with those using self-sourced planting material (Dato et al. 2021). Particularly complex and spatially interconnected planting material networks were evident in key municipalities such as Abomey-Calavi, Porto-Novo, and Dangbo, as well as across the border into Togo. These findings highlight the need to account for both social and spatial aspects in the sourcing of planting material as part of effective and sustainable plant disease management strategies.

A significant challenge identified is the widespread lack of understanding among farmers regarding BBTD and its control. More than half of respondents were unaware of BBTV, and only a minority could recognize its symptoms. This knowledge gap is a significant barrier to effective on-farm management. Compounding this challenge is the fact that early-stage symptoms of BBTD are subtle and often indistinguishable to untrained observers (Omondi et al. 2023). The variation in cultivar types, with some being more susceptible or showing different symptom expressions (Chabi et al., 2023) may further complicate disease recognition. This could explain the low rate of symptom identification and the inconsistent management practices observed. A small number of farmers believed that Banana Xanthomonas Wilt (BXM) was present in their fields. However, BXW remains confined to the African Great Lakes region and has not been detected in West Africa (Ocimati et al. 2019). Therefore, any suspected cases are likely due to lack of knowledge and misidentification, as BXW symptoms such as wilting, yellowing, or fruit rot can resemble those caused by other biotic stresses or mechanical damage (Biruma et al. 2007).

In our survey, over half of the farmers reported taking no action in response to the disease. Among those who did implement control measures, a third used decapitation of symptomatic plants and, less commonly, uprooting. Recognition of symptoms was also remarkably low, with just one tenth of farmers able to identify them in the field. This lack of awareness extends to transmission pathways, with most respondents unfamiliar with how the virus spreads, often attributing it to lack of maintenance or general pests rather than contaminated planting material or aphids. Previous studies have shown that most farmers employed a range of ad hoc, self-devised management practices, including simple cutting, uprooting (with or without manuring), chopping of the remaining pseudostem, application of ash or salty water, shifting of mats, and hot water treatment of the pseudostem (Abiola et al. 2020). The variability in these practices may reflect uneven access to or awareness of formal extension services. Nearly half of the respondents had only owned their plantations for 1–5 years, suggesting limited exposure to prior training efforts. Short cultivation cycles (≤5 years) for plantains contrast with decades-long banana cultivation, yet land tenure generally remains secure and long-term (Lescot and Ganry 2010). The findings underscore the importance of localized outreach and training programs that not only consider farmers’ experience levels but also address cultivar-specific symptom variability to improve disease management outcomes.

While men constitute the primary agricultural labour force across most activity types in banana production in sub-Saharan Africa including Benin (Ajambo et al. 2018), women are also reported to play key roles in procuring planting materials, harvesting, transporting produce for marketing, and leaf-stripping (Ahouangninou et al. 2021). In our survey, women represented 25% of respondents, a gap in sectoral involvement or visibility that was consistent across ethnic and age groups. Despite lower representation, female farmers showed comparable levels of market engagement, with the majority cultivating bananas primarily for sale. Landholding data also suggest that women were not disproportionately excluded from larger-scale production: a fifth of those managing plantations larger than 1,000 m² were women. Similarly, patterns of planting material sourcing were aligned across genders, with over 80% of both men and women relying on acquisition through neighbours. In contrast, studies from other West African countries show different patterns: in Nigeria, women more often sourced planting material from relatives, whereas men typically used old fields (Nkengla-Asi et al. 2021); in Cameroon, men were more likely than women to purchase banana planting material (Nkengla-Asi et al. 2020).

In the current study, we generated the first high-resolution national-scale map of banana cultivation in Benin. Our analysis estimates that approximately 6,663 hectares are currently under banana cultivation. This figure exceeds the most recent national estimate reported by the Food and Agriculture Organization (FAO), which indicated a harvested area of 4,282 hectares in 2023 (*FAOSTAT*. The discrepancy could be attributed to lack of ground-based agricultural statistics. For example, in the report on the latest agricultural census conducted in Benin in 2019, no data were provided on banana or plantain production (Institut National de la Statistique et de l’Analyse Économique (INSAE) 2021). Our map confirms that banana and plantain production is highly concentrated in the southern regions, with six departments accounting for over 96% of the total cultivated area. This aligns with findings from the ‘Banana Mapper Project’, a joint mapping initiative by the International Institute of Tropical Agriculture and Bioversity International to map banana-growing areas in sub-Saharan Africa (Bouwmeester et al. 2023). At the commune level, Athiémé, Toffo, Dogbo, Zè, and Lalo alone represent nearly half of the national banana production area. Our map provides the first spatially resolved, remote-sensing-based quantification of banana cultivation in Benin and offers a valuable tool for guiding agricultural policy and monitoring banana diseases (Retkute et al. 2025) or land-use change under various economic programs or future climate scenarios. For instance, a global temperature increase of 1.5–3.0 °C is projected to substantially expand plantain production areas in the Guineo-Sudanian Zone (Egbebiyi et al. 2020).

We used the model developed in Retkute & Gilligan (2025b) to predict BBTV spread and to compare management strategies. The model was parameterized using field surveillance data collected in Benin between 2018 and 2021. The survey confirmed BBTV presence in 140 of the 747 fields surveyed. No cases were detected in the northern Sudanian ecological zone, while the Guineo-Sudanian and Guineo-Congolese zones showed comparable levels of disease prevalence. This pattern aligns with the known agroecological distribution of the virus in subhumid and humid environments that support year-round banana cultivation (Kumar et al. 2011). In the complex landscape of BBTV management, mathematical modelling serves as a valuable tool to understand the spatial and temporal dynamics of disease spread. Importantly, modelling enables the evaluation of control strategies prior to implementation, helping to identify those strategies with the highest potential to reduce infection rates effectively and efficiently. By integrating spatial projections of BBTV risk with the Relative Wealth Index (RWI), our study identified regions where the risk of high disease pressure overlaps with heightened socioeconomic vulnerability. The departments of Mono, Atlantique, and Couffo were most affected, with 71.2%, 54.0%, and 45.7% of their areas falling into high BBTV risk and low-to-medium RWI categories. The communes of Athiémé, Bopa, Houéyogbé, Comè, and Grand-Popo emerged as critical nodes for disease spread, combining high BBTV risk, low RWI, and active planting material exchange. Simulations of three control strategies (decapitation, uprooting, and injection) highlighted injection-based treatment as the most effective at the national scale, reducing infected area by an average of 70%. Uprooting and decapitation were less effective, achieving reductions of 50% and 25%, respectively. However, in high-risk, low-RWI areas, uprooting proved most impactful locally, reducing risk by 70– 85%. Our results are consistent with results from field studies; for instance, the removal of infected plants reduced BBTD incidence to 2% in managed farmer fields and to 10% in experimental field plots (Omondi et al. 2020).

Overall, the results underscore the importance of tailoring BBTV control strategies not only to epidemiological conditions, but also to the social and economic contexts in which farmers operate. The prevalence of informal seed networks and low farmer awareness clearly demonstrate the need to continue education programmes on BBTV symptoms, transmission, and effective management practices. The study’s detailed mapping of production areas and identification of high-risk, vulnerable regions provides a clear roadmap for spatially informed surveillance and the deployment of clean planting material systems. While injection-based treatments show high national efficacy in simulations, their negligible adoption by farmers (0.6%) suggests barriers that need to be addressed, potentially through training and access to appropriate tools and chemicals. Cost is also likely to be a factor. The effectiveness of uprooting in high-priority areas, suggests it could be a useful and more accessible strategy for smallholder farmers in vulnerable communities. Future interventions should consider an approach integrating farmer education, promoting clean planting material systems, and implementing spatially targeted surveillance and management strategies.

## Acknowledgements

We thank all the farmers who generously shared their time, knowledge, and experiences during the surveys.

## Author contributions

R.R., M.Z.T., J.E.T., and C.A.G. planned and designed the research; M.Z.T. designed and supervised surveys; U.R.A., Y.M.V, H.A., E.A., L.D., A.M., A.E., E.H., and A.A. conducted surveys and collected data; R.R. analysed data and performed the modelling work; R.R., M.Z.T., J.E.T. and C.A.G. initiated and planned the manuscript; R.R., M.Z.T., B.A.O., U.R.A., L.K, J.E.T., and C.A.G. wrote the manuscript; all authors (R.R., M.Z.T., B.A.O., U.R.A., Y.M.V, H.A., E.A., L.D., A.M., A.E., E.H., A.A., L.K, J.E.T., and C.A.G.) have read and approved the final manuscript.

## Ethics Statement

The authors confirm that all ethical guidelines were followed in accordance with the Protection of Research Participants and the Declaration of Helsinki (1964). The research sought informed consent from the interviewees with assurances of anonymity on the purposes of the study and on how the data would be used in practice.

## References

1. Abiola, A., Adégbola, Y. P., Zandjanakou-Tachin, M., Crinot, G. F., and Biaou, G. 2024. Toward tailored interventions in plantain (*Musa paradisiaca* L.) industry: Insights from heterogeneity and constraints to plantain-based cropping systems in South-Benin. Soc. Sci. Humanit. Open 9:100895. 10.1016/j.ssaho.2024.100895.

2. Abiola, A., Zandjanakou-Tachin, M., Aoudji, K. N. A., Avocevou-Ayisso, C., and Kumar, P. L. 2020. Adoption of Roguing to Contain Banana Bunchy Top Disease in South-East Bénin: Role of Farmers’ Knowledge and Perception. Int. J. Fruit Sci. 20:720–736. 10.1080/15538362.2019.1673277.

3. Ahouangninou, C., Zandjanakou-Tachin, M., Abiola, A., Avocevou-Ayisso, C., Vodounou, M., Affokpon, A., and Fanou, A. 2021. Characterization and Typology of Banana Producing Farms in the District of Houeyogbe in Southern Benin. Curr. J. Appl. Sci. Technol. 21–32. 10.9734/cjast/2021/v40i4831640.

4. Ajambo, S., Rietveld, A. M., Nkengla, L. W., Niyongere, C., Dhed’a, D. B., Olaosebikan, D. O., Nitunga, E., Toengaho, J., Kumar, P. L., Hanna, R., Sufo Kankeu, R., and Omondi, B. A. O. 2018. Recovering banana production in bunchy top-affected areas in Sub-Saharan Africa: developing gender-responsive approaches. International Society for Horticultural Science. 10.17660/actahortic.2018.1196.27.

5. Biruma, M., Pillay, M., Tripathi, L., Blomme, G., Abele, S., Mwangi, M., Bandyopadhyay, R., Muchunguzi, P., Kassim, S., Nyine, M., Turyagyenda, L. F., and Eden-Green, S. 2007. Banana Xanthomonas wilt: a review of the disease, management strategies and future research directions.

6. Bouwmeester, H., Blomme, G., Omondi, A. B., and Ocimati, W. 2023. Banana bunchy top disease in Africa—Predicting continent-wide disease risks by combining survey data and expert knowledge. Plant Pathol. 72:1476–1490. 10.1111/ppa.13764.

7. Chabi, M. C., Dassou, A. G., Adoukonou-Sagbadja, H., Thomas, J., and Omondi, A. B. 2023. Variation in Symptom Development and Infectivity of Banana Bunchy Top Disease among Four Cultivars of Musa sp. Crops 3:158–169. 10.3390/crops3020016.

8. Chabi, M. C., Dassou, A. G., Dossou-Aminon, I., Ogouchoro, D., Aman, B. O., and Dansi, A. 2018. Banana and plantain production systems in Benin: ethnobotanical investigation, varietal diversity, pests, and implications for better production. J. Ethnobiol. Ethnomedicine 14:78. 10.1186/s13002-018-0280-1.

9. Chi, G., Fang, H., Chatterjee, S., and Blumenstock, J. E. 2022. Microestimates of wealth for all low- and middle-income countries. Proc. Natl. Acad. Sci. 119:e2113658119. 10.1073/pnas.2113658119.

10. Dato, K. M. G., Dégbègni, M. R., Atchadé, M. N., Tachin, M. Z., Hounkonnou, M. N., and Omondi, B. A. 2021. Spatial parameters associated with the risk of banana bunchy top disease in smallholder systems. PLOS ONE 16:e0260976. 10.1371/journal.pone.0260976.

11. Egbebiyi, T. S., Crespo, O., Lennard, C., Zaroug, M., Nikulin, G., Harris, I., Price, J., Forstenhäusler, N., and Warren, R. 2020. Investigating the potential impact of 1.5, 2 and 3 °C global warming levels on crop suitability and planting season over West Africa. PeerJ 8:e8851. 10.7717/peerj.8851.

12. FAOSTAT. https://www.fao.org/faostat/en/#data/QCL (accessed 16 July 2025).

13. Fink, A. H., Engel, T., Ermert, V., van der Linden, R., Schneidewind, M., Redl, R., Afiesimama, E., Thiaw, W. M., Yorke, C., Evans, M., and Janicot, S. 2017. Mean Climate and Seasonal Cycle. In Meteorology of Tropical West Africa, John Wiley & Sons, Ltd, pp. 1–39. 10.1002/9781118391297.ch1.

14. Gold, C. S., Kiggundu, A., Abera, A. M. K., and Karamura, D. 2002. SELECTION CRITERIA OF MUSA CULTIVARS THROUGH A FARMER PARTICIPATORY APPRAISAL SURVEY IN UGANDA. Exp. Agric. 38:29–38. 10.1017/S0014479702000133.

15. Hooks, C. R. R., Wright, M. G., Kabasawa, D. S., Manandhar, R., and Almeida, R. P. P. 2008. Effect of banana bunchy top virus infection on morphology and growth characteristics of banana. Ann. Appl. Biol. 153:1–9. 10.1111/j.1744-7348.2008.00233.x.

16. Institut National de la Statistique et de l’Analyse Économique (INSAE). 2021. Recensement national de l’agriculture 2019: Volume 2 – Principaux résultats du module de bas. Benin. https://instad.bj/images/docs/insae-statistiques/enquetes-recensements/RNA/Resultats-Module-base/VOLUME%202%20%20PRINCIPAUX%20RESULTATS%20DU%20MODULE%20DE%20BASE.pdf.

17. Kumar, P. L., Hanna, R., Alabi, O. J., Soko, M. M., Oben, T. T., Vangu, G. H. P., and Naidu, R. A. 2011. Banana bunchy top virus in sub-Saharan Africa: Investigations on virus distribution and diversity. Virus Res. 159:171–182. 10.1016/j.virusres.2011.04.021.

18. Lescot, T., and Ganry, J. 2010. Plantain (Musa spp.) cultivation in Africa : A brief summary of developments over the previous two decades. Proc. Int. Conf. Banana Plantain Afr. Harnessing Int. Partnersh. Increase Res. Impact Mombasa Kenya Oct. 5-9 2008. https://agritrop.cirad.fr/559683/ (accessed 16 July 2025).

19. Lokossou, B., Gnanvossou, D., Ayodeji, O., Akplogan, F., Safiore, A., Migan, D. z., Pefoura, A. m., Hanna, R., and Kumar, P. L. 2012. Occurrence of Banana bunchy top virus in banana and plantain (Musa sp.) in Benin. New Dis. Rep. 25:13–13. 10.5197/j.2044-0588.2012.025.013.

20. Madalla, N. A., Swennen, R., Brown, A., Carpentier, S., Van den Bergh, I., Crichton, R., Marimo, P., Weltzien, E., Massawe, C., Shimwela, M., Mbongo, D., Kindimba, G., Kubiriba, J., Tumuhimbise, R., Okurut, A. W., Cavicchioli, M., and Ortiz, R. 2023. Farmers’ preferences for East African highland cooking banana “Matooke” hybrids and local cultivars. Agric. Food Secur. 12:2. 10.1186/s40066-023-00407-7.

21. Minter, A., and Retkute, R. 2019. Approximate Bayesian Computation for infectious disease modelling. Epidemics 29:100368. 10.1016/j.epidem.2019.100368.

22. Nduwimana, I., Sylla, S., Xing, Y., Simbare, A., Niyongere, C., Garrett, K. A., and Bonaventure Omondi, A. 2022. Banana seed exchange networks in Burundi – Linking formal and informal systems. Outlook Agric. 51:334–348. 10.1177/00307270221103288.

23. Neuenschwander, P., and Adomou, A. C. 2017. Reconstituting a rainforest patch in southern Benin for the protection of threatened plants. Nat. Conserv. 21:57–82. 10.3897/natureconservation.21.13906.

24. Nkengla-Asi, L., Eforuoku, F., Olaosebikan, O., Adejoju Ladigbolu, T., Amah, D., Hanna, R., and Kumar, P. L. 2021. Gender Roles in Sourcing and Sharing of Banana Planting Material in Communities with and without Banana Bunchy Top Disease in Nigeria. Sustainability 13:3310. 10.3390/su13063310.

25. Nkengla-Asi, L., Omondi, A. B., Che Simo, V., Assam, E., Ngatat, S., and Boonabaana, B. 2020. Gender dynamics in banana seed systems and impact on banana bunchy top disease recovery in Cameroon. Outlook Agric. 49:235–244. 10.1177/0030727020918333.

26. Ocimati, W., Bouwmeester, H., Groot, J. C. J., Tittonell, P., Brown, D., and Blomme, G. 2019. The risk posed by Xanthomonas wilt disease of banana: Mapping of disease hotspots, fronts and vulnerable landscapes. PLOS ONE 14:e0213691. 10.1371/journal.pone.0213691.

27. Omondi, B. A., Soko, M. M., Chabi, M. C., Nduwimana, I., Adjalla, C. C., Athindehou, F., Amoussou, R. R., Dato, G. K., Tachin, M. Z., Niyongere, C., and Staver, C. 2023. Tools for the management of the banana bunchy top disease in small holder systems. 10.17660/actahortic.2023.1367.27.

28. Omondi, B. A., Soko, M. M., Nduwimana, I., Delano, R. T., Niyongere, C., Simbare, A., Kachigamba, D., and Staver, C. 2020. The effectiveness of consistent roguing in managing banana bunchy top disease in smallholder production in Africa. Plant Pathol. 69:1754–1766. 10.1111/ppa.13253.

29. Pérez-Hoyos, A., Rembold, F., Kerdiles, H., and Gallego, J. 2017. Comparison of Global Land Cover Datasets for Cropland Monitoring. Remote Sens. 9:1118. 10.3390/rs9111118.

30. Retkute, R., Crew, K. S., Thomas, J. E., and Gilligan, C. A. 2025. Detection of Banana Diseases Based on Landsat-8 Data and Machine Learning. Remote Sens. 17:2308. 10.3390/rs17132308.

31. Retkute, R., and Gilligan, C. A. 2025a. A novel two-stage parameter estimation framework integrating Approximate Bayesian Computation and Machine Learning: The ABC-RF-rejection algorithm. . 10.48550/arXiv.2507.02072.

32. Retkute, R., and Gilligan, C. A. 2025b. Developing a spatio-temporal model for banana bunchy top disease: leveraging remote sensing and survey data. Front. Plant Sci. 16. 10.3389/fpls.2025.1521620.

33. Salzmann, U., and Hoelzmann, P. 2005. The Dahomey Gap: an abrupt climatically induced rain forest fragmentation in West Africa during the late Holocene. The Holocene 15:190–199. 10.1191/0959683605hl799rp.

34. Thomas, J. Burleigh Dodds Science Publishing | Agricultural Science in Print and Online. Banana Bunchy Top Virus Achiev. Sustain. Cultiv. Banan. https://shop.bdspublishing.com/store/bds/detail/workgroup/3-190-109529 (accessed 16 July 2025).

35. Tolan, J., Yang, H.-I., Nosarzewski, B., Couairon, G., Vo, H. V., Brandt, J., Spore, J., Majumdar, S., Haziza, D., Vamaraju, J., Moutakanni, T., Bojanowski, P., Johns, T., White, B., Tiecke, T., and Couprie, C. 2024. Very high resolution canopy height maps from RGB imagery using self-supervised vision transformer and convolutional decoder trained on aerial lidar. Remote Sens. Environ. 300:113888. 10.1016/j.rse.2023.113888.

36. Trochim, W. 2007. The Research Methods Knowledge Base.

37. Vodounou, M. Y., Agoi, U., and Zandjanakou-Tachin, M. 2022. Banana Bunchy Top Disease (BBTD): Distribution, incidence and farmers’ knowledge in Benin. 32.

38. Yu, Q., You, L., Wood-Sichra, U., Ru, Y., Joglekar, A. K. B., Fritz, S., Xiong, W., Lu, M., Wu, W., and Yang, P. 2020. A cultivated planet in 2010 – Part 2: The global gridded agricultural-production maps. Earth Syst. Sci. Data 12:3545–3572. 10.5194/essd-12-3545-2020.

